# Evaluation of Autophagy in conjunctival fibroblasts

**DOI:** 10.1101/2024.01.27.575831

**Authors:** Parvaneh Mehrbod, Paola Brun, Umberto Rosani, Andrea Leonardi, Saeid Ghavami

**Author notes:** Corresponding Author, Saeid Ghavami, PhD, Department of Human Anatomy and Cell Science, University of Manitoba College of Medicine, Winnipeg, MB R3E 0V9, Canada. These authors have co-first authorship (Parvaneh Mehrbod, Paola Brun). These authors have co-senior authorship (Andrea Leonardi, Saeid Ghavami).

## Abstract

Vernal keratoconjunctivitis (VKC) is a serious eye allergy characterized by poorly understood pathogenic mechanisms and a lack of effective treatments. Autophagy, a process involved in both triggering and suppressing immune and inflammatory responses, plays a role in VKC’s pathophysiology. Understanding autophagy’s involvement in VKC could lead to new treatment possibilities, such as utilizing specific topical substances to induce or inhibit autophagy and prevent severe complications of this eye condition. In our current protocol, we present a robust methodology established in our laboratory for studying autophagy in primary conjunctival fibroblasts. We assess autophagy through techniques like immunocytochemistry, immunoblotting, and qPCR.

## 1 Introduction

Macroautophagy, also known as autophagy, is a meticulously controlled catabolic process preserved throughout evolution, functioning at baseline levels in the majority of cells to maintain cellular equilibrium by breaking down surplus proteins and eliminating impaired organelles through lysosomal degradation [1-4]. Yet, it can swiftly escalate in reaction to various metabolic stresses such as nutrient deficiency, oxygen deprivation, or heightened metabolic demands associated with rapid cell growth, all aimed at preserving cellular balance [5-8]. Recent advancements reveal a nuanced and multifaceted relationship between the autophagy pathway and immunity [9, 10].

The autophagy pathway plays a role in both triggering and dampening immune and inflammatory reactions. Conversely, chemokines and cytokines can either stimulate or hinder the autophagy pathway in mammalian cells [11-14]. MyD88 and TRIF (Toll-like receptor adaptors) along with HGB1 disrupt the interaction between BECLIN-1 and BCL2 by binding to BECLIN-1 and displacing BCL2.[15, 16]. Additional evidence regarding the regulation of autophagy and immune signaling molecules was presented through genome-wide siRNA screening [17]. In this analysis, 219 genes were identified that inhibit basal autophagy, operating independently of the mammalian target of rapamycin complex 1 *(mTORC1*). These genes encompassed cytokines (*CLCF1, LIF, IGF1, FGF2*), a chemokine (*SDF1*, also recognized as *CXCL12*), and cellular signaling molecules influenced by cytokines (*STAT3*) [17]. Considering the broad role of autophagy in maintaining cellular balance and modulating immune and inflammatory signaling, alterations in autophagy levels induced by cytokines in immune cells could have a significant impact on immunity and inflammation [9].

Studies have demonstrated the essential role of autophagy in the secretion of bronchial epithelial mucus in a Th2 model of asthma [18]. Autophagy might also participate in airway remodeling, presenting a potentially innovative focus in treating steroid-resistant severe asthma, indicating its involvement in allergic conditions [14, 18].

Inflammation and autophagy are pivotal biological processes engaged in various physiological and pathological scenarios. Pro-inflammatory cytokines exert autophagy-mediated effects across diverse inflammatory conditions. The interplay between autophagy and inflammation represents a fundamental mechanism regulating the function of both the innate and adaptive immune systems [19]. Autophagy has the capacity to both regulate and be influenced by a broad spectrum of pro-inflammatory cytokines [20, 21]. These interactions have the potential to either ameliorate or exacerbate inflammatory states. Consequently, the interplay between autophagy and inflammatory cytokines holds significant importance and serves as a primary focus of research in modulating autophagy and immunity [20].

Vernal keratoconjunctivitis (VKC) is a serious and uncommon chronic eye allergy that predominantly impacts children and teenagers residing in warm regions, significantly disrupting their daily lives and overall well-being [22]. The condition is marked by severe ocular symptoms including itching, sensitivity to light, a feeling of foreign objects in the eye, redness of the conjunctiva, and excessive mucus production. These symptoms are often aggravated during or after outdoor activities. Additionally, patients may develop large papillae on the upper eyelid or inflammation and nodules near the cornea. This discomfort not only significantly impacts the quality of life of patients and their caregivers but also poses risks of complications such as cataracts and glaucoma due to long-term corticosteroid treatment [23].

VKC has been recognized as an allergic reaction mediated by both IgE and T cells, wherein the CD4+ T helper (Th)2 inflammatory pattern, prompted by IL-4, plays a significant role. This pattern involves the secretion of IL-4, IL-5, and IL-13 by cells and has been shown to influence different levels of corneal involvement and tissue restructuring [24-26].

Similar to other atopic conditions like atopic dermatitis and asthma, the precise trigger for VKC often remains unknown in the majority of cases. A range of nonspecific factors, including warm climates, sunlight, dry air, oxidative stress, and potential panallergens, can worsen or sustain ocular inflammation. In VKC patients, there is an increase in various cytokines (including Th2-type and Th1-type), chemokines, growth factors, and enzymes [27, 28], suggesting VKC is associated with unique patterns of inflammation in the conjunctiva.

Studies have demonstrated that the activities and characteristics of conjunctival fibroblasts and epithelial cells can be altered in laboratory settings by various mediators implicated in ocular allergic reactions, including histamine, IL-1, IL-4, and TNF-α [28-33]. These cytokines also play a role in triggering autophagy, underscoring the interconnectedness between autophagy, inflammatory agents, and the development of VKC.

Histamine, a biogenic amine with diverse biological roles, triggers an elevation in the ratio of LC3II/LC3I and stimulates the production of intracellular reactive oxygen species (ROS), consequently inducing autophagy that is stress-induced within the endoplasmic reticulum (ER) [34]. TNF-α, a multifaceted cytokine known for its modulation of pro-inflammatory reactions, is implicated in interfering with the process of autophagy IL-1β binds to IL-1 receptor 1, initiating inflammatory cascades by stimulating the production of IL-1α and IL-23. This, in turn, prompts the formation of autophagosomes, which may induce autophagy as a component of a negative feedback mechanism to mitigate the inflammatory response [35]. IL-4, a crucial Th2 cytokine, serves as a key effector in the immune response. Harris et al. demonstrated that unlike pro-inflammatory cytokines, the secretion of anti-inflammatory cytokines like IL-4 inhibits the induction of autophagy by activating mTOR [36].

Hence, in this chapter, we introduced a novel methodology for assessing autophagy in conjunctivitis fibroblasts in response to inflammatory mediators. We employed established techniques including immunohistochemistry, qPCR, and Western Blotting (WB).

## 2 Materials

### 2.1. Conjunctival biopsies

1. Optimal cutting temperature (OCT) compound (Histo-Line Laboratories, Pantigliate, MI, Italy)

### 2.2. Immunohistochemistry (IHC)

1. Superfrost plus slides (Thermo scientific, Waltham, MA, USA)
2. Formaldehyde (VWR International, Milan, Italy)
3. TRIZMA (2-Amino-2-(hydroxymethyl)-1,3-propanediol, tris (hydroxyamino methane) maleate buffer 0.05 M, pH 7,6 (Sigma, St. Louis, MO, USA)
4. Normal horse serum (Vector Laboratories Inc, Burlingame, CA, USA – S 2000)
5. Normal rabbit serum (Vector Laboratories Inc, Burlingame, CA, USA – S 5000)
6. Normal goat serum (Vector Laboratories Inc, Burlingame, CA, USA – S 1000)
7. Secondary antibody anti-mouse made in horse (Vector Laboratories Inc, Burlingame, CA, USA - BA 2000
8. Secondary antibodies anti-rabbit made in goat (Vector Laboratories Inc, Burlingame, CA, USA - BA 1000)
9. Secondary antibodies anti-goat made in rabbit(Vector Laboratories Inc, Burlingame, CA, USA - BA 5000)
10. Rabbit, goat or mouse non-specific immunoglobulins (Santa Cruz Biotechnologies, Santa Cruz, CA, USA)
11. ABC kit HRP Elite (Vectstain, Vector Laboratories Inc, Burlingame, CA, USA - PK6100)
12. Mayer Heantoxylin (AppliChem, Darmstadt, Germany)

### 2.3. Cell cultures and tretments

1. Type I collagenase (Worthington Biochemical, Lakewood, NJ, USA)
2. Wong-Kilbourne derived Chang (ChWK) conjunctival epithelial cells (American Type Culture Collection -ATCC, Manassas, VA, USA)
3. Human monocytes U937 (Thermo Scientific (Wilmington, DE, USA)
4. Dulbecco’s Modified Eagle’s Medium (DMEM, Gibco; Grand Island, NY. USA)
5. RPMI 1640 medium (Gibco; Grand Island, NY, USA)
6. 10% Fetal Bovine Serum (FBS) (Gibco; Grand Island, NY, USA).
7. Penicillin/streptomycin (P/S, Euroclone, Milan, Italy),
8. Phorbol-12-myristate-134 acetate (PMA, Sigma, St. Louis, MO, USA)
9. Lipopolysaccharide (LPS, Sigma, St. Louis, MO, USA)
10. Histamine (Sigma, St. Louis, MO, USA), IL-1β, IL-4 and TNF-α (Peprotech, London, UK)
11. Insulin, Transferrin, Selenium supplement (ITS, GIBCO, Grand Island, NY, USA)
12. Chloroquine (CQ, Sigma, St. Louis, MO, USA)

### 2.4. Single and double staining immunofluorescence

1. Coverslips (Thermo scientific, Waltham, MA, USA)
2. Human primary conjunctival fibroblasts obtained as described above
3. TNF-α (Peprotech, London, UK)
4. DMEM medium (Gibco; Grand Island, NY. USA)
5. Insulin, Transferrin, Selenium supplement (ITS, GIBCO, Grand Island, NY, USA)
6. Formaldehyde (VWR International, Milan, Italy)
7. PBS, Triton X-100 and BSA (Sigma, St. Louis, MO, USA)
8. Anti-goat IgG conjugated with Alexa Fluor 488 (Invitrogen, Thermo scientific, Waltham, MA, USA)
9. Anti-rabbit IgG conjugated with Alexa Fluor 647 (Invitrogen, Thermo scientific, Waltham, MA, USA)
10. Hoechst (Sigma, St. Louis, MO, USA)
11. Glycerol and DABCO (Sigma, St. Louis, MO, USA)

### 2.5. RNA isolation and qPCR analysis

1. TRIzol (Life Technologies, Carlsbad, CA, USA)
2. Oligo(dT) (Life Technologies, Carlsbad, CA, USA)
3. SuperScript II Reverse Transcriptase 200 U (Life Technologies, Carlsbad, CA, USA)
4. SYBR Green I dye (Roche, Mannheim, Germany)

### 2.6. Western Blotting

1. Triton X-100, deoxycholic acid, EDTA, Tween 20 and PBS (Sigma, St. Louis, MO, USA)
2. Bradford reagent (Serva, Heidelberg, Germany)
3. Acrylamide solution (AppliChem, Darmstadt, Germany)
4. Tris pH 6.8, Glycerol, sodium dodecyl sulphate (SDS), β-mercaptoethanol and bromophenol blue, glycine and Methanol (Sigma, St. Louis, MO, USA)
5. Dry milk (AppliChem, Darmstadt, Germany)
6. Horseradish Peroxidase (HRP)-conjugated anti-mouse IgG (GE Healthcare, Buckinghamshire, UK)
7. Horseradish Peroxidase (HRP)-conjugated anti-rabbit IgG and anti-goat IgG (R&D Systems, Minneapolis, MN, USA)
8. ECL Plus (GE Healthcare, Buckinghamshire, UK)
9. Anti-β-actin antibody (Sigma, St. Louis, MO, USA)

## 3 Methods

### 3.1. Conjunctival biopsies

1. Prepare conjunctival tissues from active VKC patients and healthy age-matched control subjects who underwent eyelid surgery. (**see Note 1 and Note 2**)
2. Snap freeze upper tarsal conjunctival biopsies with optimal cutting temperature (OCT) compound (Histo-Line Laboratories, Pantigliate, MI, Italy) in liquid nitrogen and maintaine at -80°C until use for the immunohistochemistry (data not shown) and qPCR analysis. (**see Note 3**)

### 3.2. Immunohistochemistry (IHC)

1. Stain two sections from each biopsy sample applying immunohistochemical methods with a panel of antibodies specific for some autophagic components (Table 1).
2. Prepare serial 5 μm-thick cryosections and fix them in 4% buffered formaldehyde and incubate with 0.3% hydrogen peroxide for 30 min. (**see Note 4**)
3. Block the samples with appropriate serum (1:20) in TRIZMA (2-Amino-2-(hydroxymethyl)-1,3-propanediol, tris (hydroxyamino methane, Sigma, St. Louis, MO, USA), and maleate 0.05 M, pH 7.6.
4. Incubate the sections with adequate dilution of the primary antibody in TRIZMA maleate for 1 hr at room temperature in a humid chamber.
5. Demonstrate antibody binding using secondary antibodies anti-mouse (Vector, BA 2000), anti-rabbit (Vector BA 1000) or anti-goat (Vector, BA 5000) followed by ABC kit HRP Elite, PK6100 (Vectstain) and diaminobenzidine substrate (brown color) (Sigma, St. Louis, MO, USA).
6. Counterstain the slides with heantoxylin and than rinse with tap water.
7. Perform negative controls using normal non-specific goat, mouse or rabbit immunoglobulins (Santa Cruz Biotechnologies, Santa Cruz, CA, USA).
8. Observe with a light microscope (Leica Axioplan, Wetzlar, Germany) at 400x magnification.
9. Evaluate the positive reaction for all the studied antigens in the epithelium and sub-epithelial stroma of conjunctival tissues using a 0-3 score by two independent investigators. (**see Note 5**)

### 3.3. Cell cultures and tretments

#### 3.3.1. Methods per cell cultures and inflammatory conditioned medium (CM) production

1. Obtain primary conjunctival fibroblasts after digestion of VKC tarsal biopsies with 20 U/ml type I collagenase (Worthington Biochemical) at 37°C for 12 hr.
2. Prepare Wong-Kilbourne derived Chang (ChWK) conjunctival epithelial cells from the American Type Culture Collection (ATCC; Manassas, VA, USA) as control cell lines.
3. Obtaihe human monocytes U937 from Thermo Scientific (Wilmington, DE, USA) for the production of the CM.
4. Grow the conjunctival cells under the standard cell culture practices in complete Dulbecco’s modified Eagle’s medium (DMEM) supplemented with 10% fetal bovine serum (FBS), 1% (v/v) penicillin/streptomycin (P/S), 2 mM glutamine at 37°C, in a 5% humidified CO_2_ atmosphere.
5. Culture U937 cells in suspension in complete RPMI 1640 medium (Gibco; Grand Island, NY) and sub-culture twice weekly by dilution using a seeding density of 10^6^ cells/ml.
6. Differentiate U937 monocytes to macrophages by addition of 20 ng/ml of phorbol-12-myristate-134 acetate (PMA) into the cell culture medium for 48 hr and subsequently incubate for 1 hr with 1 μg/ml lipopolysaccharide (LPS).
7. Examine differentiation of monocytes to macrophages under an inverted phase-contrast microscope and also assess the mRNA expression of CD68 (the macrophage differentiation marker) by quantitative real-time PCR (qPCR).
8. Wash the cells and culture in complete DMEM for 24 hr to produce the inflammatory conditioned medium (CM).
9. Subsequently withdraw, centrifuge, filter and store at -80°C.

#### 3.3.2. Methods for conjunctival cells treatments

1. Seed VKC conjunctival fibroblasts in 6-well culture dishes and treat for 24 hr with the U937 culture medium (CM). (**see Note 6**)
2. Prior to each treatment, starve the cell cultures with DMEM in the presence of 1% insulin, transferrin, and selenium supplement (ITS) for 24 hr, to make them quiescent. (**see Note 7**)
3. Replace the medium by complete DMEM medium and detach the cells at 4, 10 and 24 hr and freeze at -80°C until lysis for protein and mRNA extraction to analyze the expression of autophagic markers. (**see Note 8**)
4. In another series of experiments seed both conjunctival cell types in 6-well culture dishes and treat with 10 ng/ml of histamine, 1 ng/ml of human recombinant IL-1β, 10 ng/ml IL-4, or 10 ng/ml TNF-α. (**see Note 9**)
5. In some experiments, expose TNF-α-treated fibroblasts in 1% ITS to 100 μM CQ for 4, 10 and 24 hr and subsequently analyze LC3B expression by Western Blotting. (**see Note 10**)

### 3.4. Single and double staining immunofluorescence

1. Seed conjunctival fibroblasts at a concentration of 5000 cell/cm^2^ coverslips at the bottom of 24-well culture dishes and cultivate in the appropriate medium with 10% FBS.
2. Allow the cells to grow to 50% confluency and treat them with 10 ng/ml TNF-α in DMEM with 1% ITS for 4, 10 and 24 hr.
3. Fix the cells in 4% formaldehyde and quench with 100 mM glycine in PBS and permeabilize in 0.25% Triton X-100.
4. Block the cells with 3% bovine serum albumin (BSA) in phosphate buffered saline (PBS) fo 1 hr at -20°C.
5. Incubate the slides for the single reaction with the primary anti-LC3B antibody, and for the double reaction with the rabbit anti-LAMP1 for 1 hr and subsequently with the goat anti-Cathepsin D antibody overnight at 4°C.
6. Wash the cells with PBS, then incubate them with the appropriate secondary antibodies (anti-goat IgG conjugated with Alexa Fluor 488; anti-rabbit IgG conjugated with Alexa Fluor 647) diluted at 1:200 in PBS.
7. Counterstain the nuclei with Hoechst (Sigma) for 5 min and mount the slides using 90% glycerol and 5% DABCO.
8. Examine the antibody reactivity with fluorescence microscope (Zeiss, Heidelberg, Germany) by two independent observers. (**see Note 11**)

### 3.5. RNA isolation and qPCR analysis

1. Extract total RNA from conjunctival tissues, conjunctival cell cultures or U937 cell cultures using TRIzol (Life Technologies, Carlsbad, CA, USA) according to the manufacturer’s protocol.
2. Quantify the RNA concentration of each sample using NanoDrop 2000c Spectrophotometer (Thermo Scientific, Wilmington, DE, USA).
3. Use the primers pairs (Table 2). (**see Note 12**)
4. Retro-transcribe 1 μg of total RNA into cDNA using oligo(dT)_12-18_, 25 ng of primers and 1 μl of SuperScript II Reverse Transcriptase 200 U (Life Technologies).
5. Perform the qPCR analysis with real-time PCR system (Rotor Gene RG-3000A, Corbett Research) using SYBR Green I dye (Roche, Mannheim, Germany) and combine sense and antisense primers at 300 nM final concentration.
6. Use the following cycling conditions: initial denaturation step of 10 min at 95°C, 45 cycles of amplification consisting of 15 s at 95°C, 30 s at 60°C, and 30 s at 72°C.
7. Evaluate genes expression with δδCt method using peptidylprolylisomerase A (PPIA) as reference gene. (**see Note 13**)

### 3.6. Western Blotting

1. Analyze conjunctival fibroblasts and epithelial cells by Western Blotting for the presence of LC3B, Beclin-1,Cathepsin D and p62 (Table 1).
2. Treat cell pellets with RIPA lysis buffer (1% v/v Triton X-100, 0.5% w/v deoxycholic acid, 10 mM EDTA in PBS, Sigma) supplemented with protease inhibitor cocktail for 45 min at 4°C.
3. Remove the particulate material by centrifugation (12,000 g for 10 min, at 4°C), collect the supernatants, and determine protein concentration using the Bradford Assay.
4. Add the lysates (20 μg) to the sample loading buffer (62.5 mM Tris pH 6.8, 10% v/v glycerol, 2% w/v sodium dodecyl sulphate, 5% v/v β-mercaptoethanol, and 0.1% w/v bromophenol blue), denature at 98°C for 5 min and separate by 10% sodium dodecyl sulphate-polyacrylamide gel electrophoresis (SDS-PAGE).
5. Transfer the proteins to the nitrocellulose membrane overnight at 4°C with the constant current of 100 mA, in blotting buffer (25 mM Tris, 192 mM Glycine and 20% Methanol).
6. Block non-specific binding sites incubating the nitrocellulose membrane for 1 hr at 23ºC in 5% w/v non-fat dry milk in 20 mM Tris [pH 7.6], 150 mM NaCl (TBS).
7. Incubate the nitrocellulose membrane with anti-LC3B, LC3A, Beclin-1, Cathepsin D or p62 appropriately diluted antibodies in TBS, 0.1% Tween-20 (TBST) for 2 hr at 23ºC.
8. Wash the membrane three times for 10 min in TBST (Tris-Buffered Saline and Tween 20, Sigma, St. Louis, MO, USA).
9. Incubate the membrane with HRP (Horseradish Peroxidase)-conjugated antibody (anti-mouse IgG, anti-rabbit IgG or anti-goat IgG) diluted in TBST, for 1 hr at 23ºC.
10. Reveal the antibody reaction by chemiluminescence using ECL Plus (GE Healthcare).
11. Incubate the blots sequentially at 23ºC for 2 hr with monoclonal anti-β-actin antibody (Sigma) diluted at 1:4000 and process as described above.
12. Normalize the densitometry values of positive bands with the corresponding value of β-actin.
13. For the statistical analysis, compare the groups using Student’s unpaired *t*-test. Compare the protein levels between VKC and control biopsies using the non-parametric Mann-Whitney U-test. (**see Note 14 and Note 15**)

### 3.7. Interpretation of results

For autophagy evaluation, the autophagy proteins are first evaluated in conjunctivitis tissue using IHC. An example of immunohistochemical analysis for BECLIN in VKC biopsy was shown in Figure 1.

**Table 1.** Autophagy markers.

| Target | Supplier | Catalogue number | Source | Methods |
| --- | --- | --- | --- | --- |
| **MAP1LC3A** | abcam | ab52628 | rabbit | IHC |
| **LC3B** | abcam | ab64782 | rabbit | IHC |
| **LC3B** | Proteintech | 18725-1-AP | rabbit | IF |
| **LC3B** | Sigma | L7543 | rabbit | WB |
| **Beclin-1** | Cell signalling Tech. | #3495P | rabbit | IHC, WB |
| **Cathepsin D** | Santa Cruz Biotech. | Sc-6486 | goat | IHC, WB, IF |
| **P62** | Millipore | MABC32 | mouse | WB |
| **LAMP 1** | Sino Biological Inc | 11215-R107 | rabbit | IHC, IF |

**Table 2.**
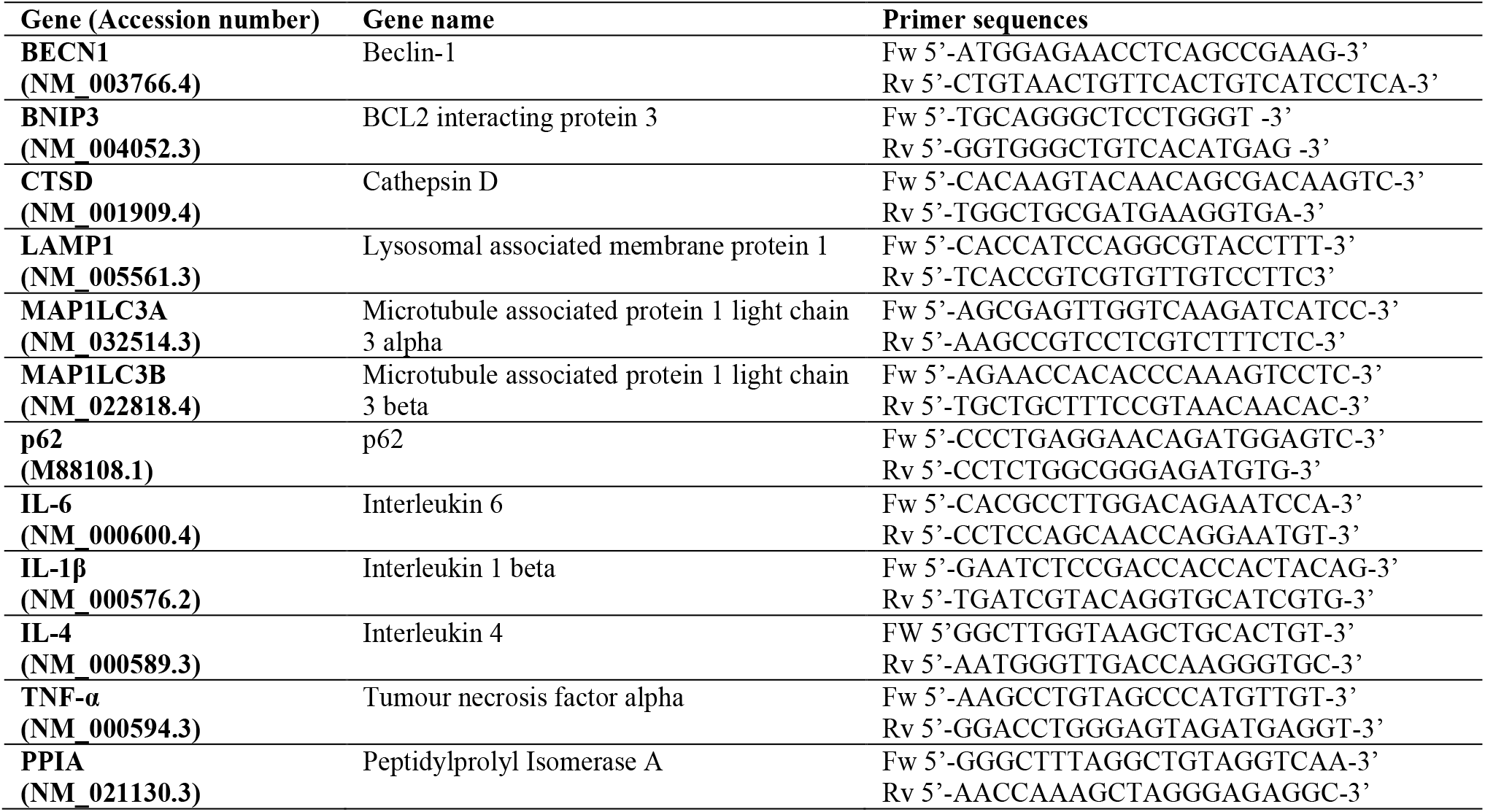
Genes to investigate and primers to use.

**Figure 1.**
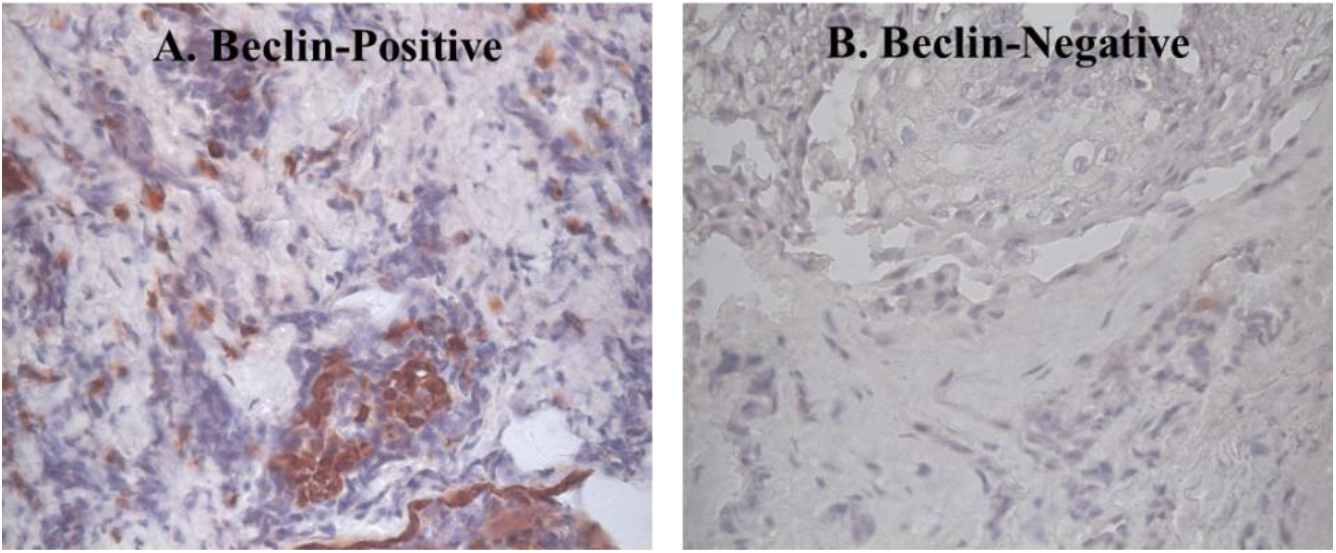
Immunohistochemical analysis for Beclin in human vernal keratoconjunctivitis (VKC) biopsy. The biopsy samples were treated with the anti-Beclin antibody. A) the positive reaction was obtained in the stroma of conjunctival tissue incubated with the anti-beclin antibody for 1 hour. B) Negative control was performed using normal nonspecific rabbit immunoglobulins.

Following that, the positive cells are cultured and modulated by diverse mediators involved in the ocular allergic reaction. Conducting qPCR, the results will show increase in the autophagy-related genes that guide us for the next step; IHC. Then, the colocalization of autophagosome and lysosome markers in IHC will guide us for another confirmation by WB. By switching to WB, LC3 lipidation, p62 degradation and other related markers are evaluated. Application of CQ for confirmation of autophagy flux will be the final step. Figure 2 shows an example of WB analysis.

**Figure 2.**
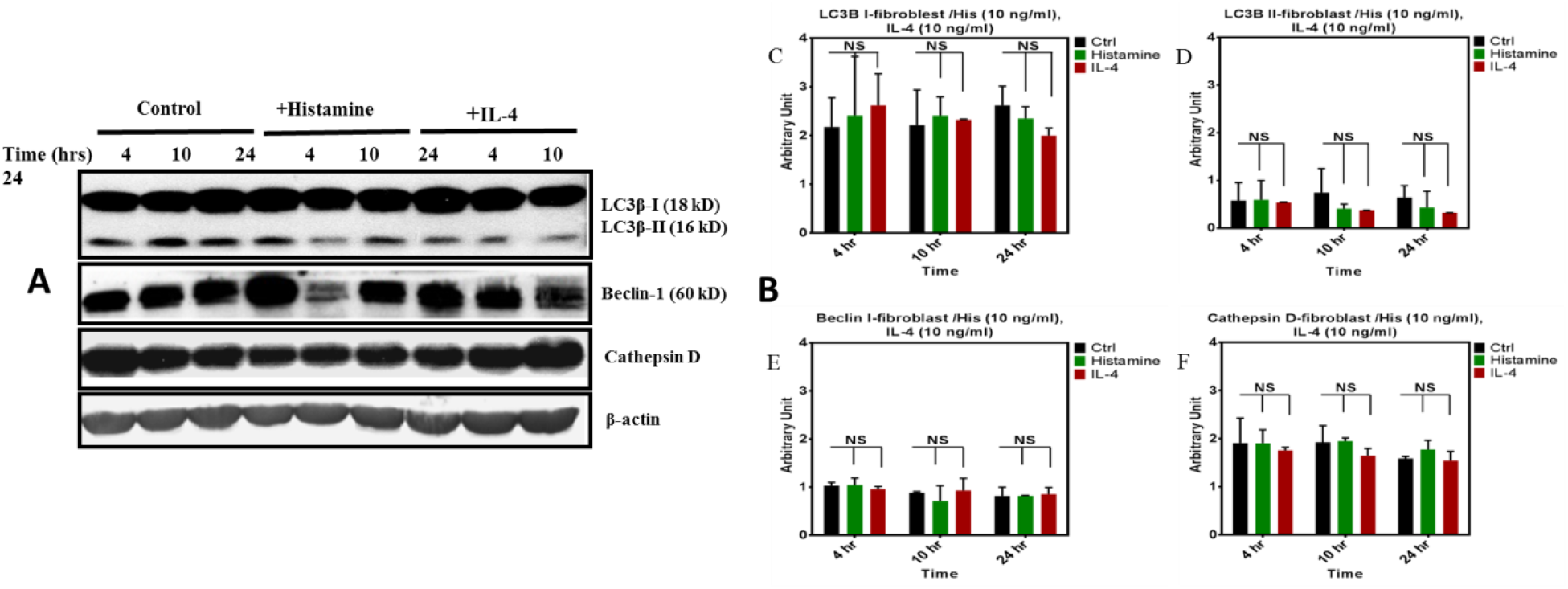
Western Blot analysis of autophagic markers expression in conjunctival fibroblasts stimulated by Histamine and IL4. **A**) Primary human conjunctival fibroblast cultures were stimulated by histamine or IL-4 and then the expression of LC3BI and II, BECLIN and Catephsin D were evaluated by WB analysis at 4, 12 and 24 hr after stimulation. B) Densitometric values of positive bands were normalized with the corresponding value of β-actin. The results demonstrated that *in vitro* expression of LC3B, BECLIN and Catephsin D was not changed.

A schematic of the methods used is shown in Figure 3.

**Figure 3.**
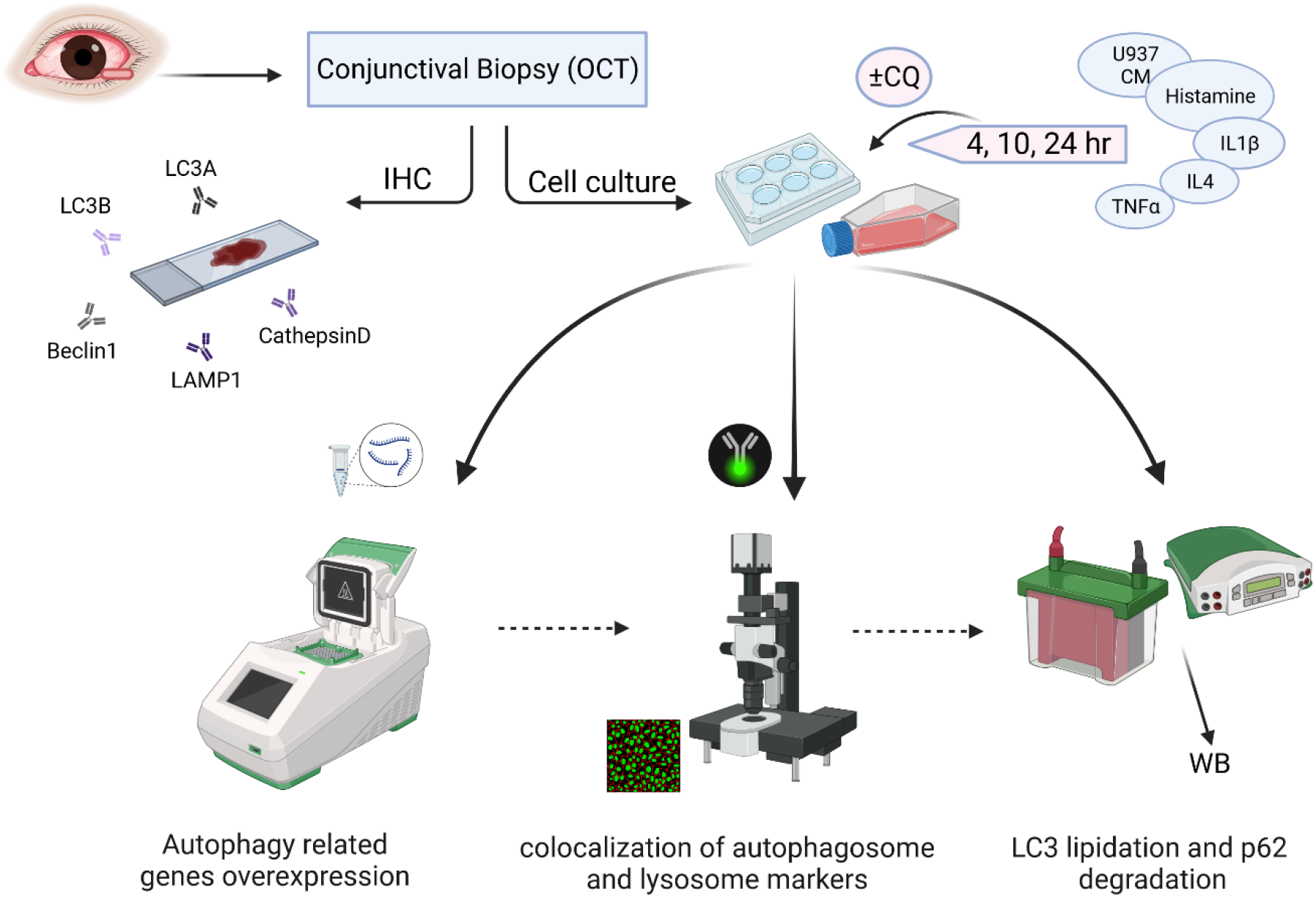
The schematic of the methods used for this methodology. (Created with BioRender.com)

## 4. Notes

1. None of the control subjects should use contact lenses or have inflammatory signs and symptoms or history of allergy.
2. Diagnosis of VKC should be based on the typical clinical history and evaluation of the signs and symptoms. All VKC patients must be free of topical mast cell stabilizers and/or anti-histamines for at least three days and free of topical corticosteroids for at least 7 days before tissue samples collection. VKC patients may be positive to serum specific IgE for at least one aeroallergen, such as bluegrass, mites, parietariae, compositae and/or tree pollens.
3. A written informed consent must be obtained from all the subjects or their parents before obtaining tissue specimens. The research study should be approved by the Institutional Review Board and Local Ethical Committee, and adhered to the tenets of the Declaration of Helsinki.
4. Incubation with 0.3% hydrogen peroxide for 30 min is necessary to inhibit endogenous peroxidase activity (IHC).
5. 0-3 score is as follow: (from 0 = absence of immunostaining to 3 = extensive intense immunostaining).
6. Autophagic proteins expression is analyzed in ChWK and conjunctival fibroblasts stimulated with activated U937 culture medium (CM) or with various pro-inflammatory molecules.
7. Making cells quiescent ensures that all the cells are synchronized in the same phase of the cell cycle and when stimulated their responses would be comparable.
8. The experiment is repeated three times to make sure of the accuracy of the results.
9. The preliminary unpublished data in this regard showed that these concentrations of the cytokines yielded the best results for the purposes of this study.
10. The reason to expose the cells to chloroquine (CQ) is that CQ treatment inhibits the autophagic flux [37] and is used as control treatment.
11. The immunelouresece protocol has been performed according to our established method which has been described in our previous investigations [38, 39].
12. The primers pairs (Table 2) were designed by Primer3 from a DNA sequence program and synthesized.
13. The cDNA samples must be analyzed in triplicate.
14. For the statistical significance, differences are indicated at *P*<0.05 (*), *P*<0.01 (**) and *P*<0.001 (***).
15. Immunoblotting has been performed according to our established protocol [40, 41].

## Acknowledgement

SG and AL conceptualized and designed the study. PB performed the experiments. SG and PB and PM analyzed the data. PM, and PB wrote the manuscript. UR, PM, PB, AL, and, SG reviewed and edited the manuscript. AL funded and SG supervised the experiments

